# The patterned assembly and stepwise Vps4-mediated disassembly of composite ESCRT-III polymers drives archaeal cell division

**DOI:** 10.1101/2022.09.16.508273

**Authors:** Fredrik Hurtig, Thomas C.Q. Burgers, Alice Cezanne, Xiuyun Jiang, Frank Nico Mol, Jovan Traparić, Gabriel Tarrason-Risa, Andre Arashiro Pulschen, Lena Harker-Kirschneck, Anđela Šarić, Rifka Vlijm, Buzz Baum

## Abstract

ESCRT-III family proteins form composite polymers that deform and cut membrane tubes in the context of a wide range of cell biological processes across the tree of life. In reconstituted systems sequential changes in the composition of ESCRT-III polymers induced by the AAA ATPase Vps4 have been shown to remodel membranes. However, it is not known how composite ESCRT-III polymers are organised and remodelled in space and time in cells. Here, taking advantage of the relative simplicity of the ESCRT-III-dependent division system in *Sulfolobus acidocaldarius*, one of the closest experimentally tractable prokaryotic relative of eukaryotes, we use super-resolution microscopy and computational modelling to show how CdvB/CdvB1/CdvB2 proteins form a precisely patterned composite ESCRT-III division ring which undergoes stepwise Vps4-dependent disassembly and contracts to cut cells into two. These observations lead us to suggest sequential changes in a patterned composite polymer as a general mechanism of ESCRT-III-dependent membrane remodelling.

## Introduction

ESCRT-III proteins form polymers that play a critical role in membrane remodelling across the tree of life. In bacteria, ESCRT-III proteins, variably termed PspA/Vipp1/IMO30, have been shown to function in membrane repair and biogenesis (*1, 2*). In archaea and eukaryotes, ESCRT-III proteins form polymers that function together with the AAA-ATPase Vps4 to cut membrane tubes from the inside to execute a wide variety of biological processes from membrane repair to compartment sealing, vesicle formation, and abscission (*3–6*). Although the precise mechanism by which ESCRT-III proteins and Vps4 remodel and cut membranes remains an unresolved question in the field, purified ESCRT-III proteins have been shown to form composite polymers that assume a wide variety of forms, from flat spirals, to cones and helices (*7–12*). In addition, reconstitution experiments have shown that ESCRT-III co-polymers can undergo stepwise changes in their composition in the presence of ATP and Vps4, which can drive changes in membrane curvature that lead to membrane tube formation and, ultimately, membrane scission (*13–17*). Furthermore, physical computational models of the process show that sequential changes in polymer structure are likely sufficient to drive membrane remodelling and scission (*18, 19*). In addition, a recent theoretical analysis (*18*) has suggested that differences in the structure of different ESCRT-III polymers and their affinity for membranes, when aided by the disassemblase activity of the AAA-ATPase, Vps4, are sufficient to generate the ordered changes in copolymer composition required for membrane deformation.

Currently, however, there is little evidence for stepwise changes in polymer organisation driving membrane remodelling and scission *in vivo*. An exception to this is the case of cell division in *Sulfolobus acidocaldarius*, one of the closest experimentally tractable archaeal relatives of eukaryotes (*20, 21*). In these archaea, as is the case for several other TACK and Asgard archaea (*22–25*), cell division appears to depend on a set of ESCRT-III proteins working together with Vps4 (also termed CdvC) (*24, 26*). Because the ~1.25 μm diameter *S. acidocaldarius* division rings assemble and constrict with precise timing during the cell cycle, these cells provide an excellent model system in which to dissect the events involved in ESCRT-III/Vps4-dependent membrane remodelling. Using this system, we recently showed that proteasomal degradation of CdvB, one of the three main ESCRT-III proteins triggers the transition from medial ring assembly to cytokinetic ring constriction and abscission (*27*). However, technical limitations prevented us from observing subcellular organisation of the different ESCRT-III polymers. Here, using a combination of ~30 nm resolution Stimulated Emission Depletion (STED) microscopy, live imaging, together with computational modelling we have been able to determine the subcellular organisation of the ESCRT-III polymers during division and to explore the role of Vps4 in this process. By studying division in control cells and in cells expressing a dominant negative version of Vps4 we show that: i) CdvB, CdvB1 and CdvB2 polymers exhibit distinct behaviours and preferred curvatures, ii) CdvB, CdvB1 and CdvB2 assemble at spatially distinct positions relative to one another at the cell midzone to generate a pre-patterned composite division ring, iii) CdvB1 and CdvB2 polymers maintain their relative positions as the rings constrict, and iv) membrane scission depends on disassembly of polymeric CdvB2 from the cytokinetic bridge. Taken together, these data lead us to propose a general framework by which to understand ESCRT-III/Vps4 function during archaeal cell division and other instances of ESCRT-III-dependent membrane remodelling in both archaea and eukaryotes.

## Results

To investigate the role of Vps4 in the regulation of ESCRT-III-mediated cell division in *S. acidocaldarius*, we generated an arabinose-inducible expression plasmid carrying Vps4 with a point mutation in the Walker B motif **(Fig. 1A),** hereafter called Vps4WkB. This point mutation prevents AAA-ATPases like Vps4 from hydrolysing bound ATP and can therefore interfere with the activity of endogenous Vps4 complexes when over-expressed (*24, 28*). The induction of Vps4WkB expression in *S. acidocaldarius* led to a significant increase in the population of large cells with a DNA content of greater than 2N **(Fig. 1B, S1A)**, as previously reported (*24*), and to an increase in the percentage of cells containing elevated levels of the ESCRT-III proteins, CdvB, CdvB1 and CdvB2 **(Fig. S1B)**, and Vps4WkB **(Fig. S1C)**. In these cultures, the majority of cells with a >2N DNA content expressed high levels of CdvB, CdvB1 and CdvB2 **(Fig. 1C)**. At the same time, we also observed a corresponding loss of newly divided G1 cells **(Fig. S1D)**, identified as a population of cells with a 1N DNA content and high levels of CdvB1 and CdvB2. This is consistent with the hypothesis that the expression of Vps4WkB prevents ESCRT-III polymer disassembly and cell division. Importantly, while a similar pattern of ESCRT-III protein accumulation and division arrest has been reported to occur following the treatment of cells with a proteasomal inhibitor (*27*), the impact of the two treatments on cell cycle progression is different. Cells treated with the proteasomal inhibitor Bortezomib are both unable to divide and unable to initiate the next round of DNA replication (*27*), whereas cells prevented from dividing as a result of Vps4WkB expression undergo multiple additional rounds of DNA replication, leading to the steady accumulation of cells with an ever-larger DNA content **(Fig. S1E)**. As a further test of this finding, we expressed the Vps4WkB plasmid in a cell line carrying a thymidine kinase homologue (hereafter called STK). This makes it possible to visualise new DNA synthesis through the incorporation of the thymidine analogue EdU by fluorescent microscopy (*29*). In control cells, EdU labelled DNA was never observed in cells undergoing DNA segregation or cell division. Instead, EdU was only incorporated into the DNA of small newly divided cells (identified by the presence of high levels of cytoplasmic CdvB1 content), as expected based on the fact that wildtype cells initiate DNA replication shortly after entering G1 **(Fig. 1D)** (*27, 30*). By contrast, when cell division was inhibited by the expression of Vps4WkB, EdU incorporation was detected in dumbbell shaped cells and in fully constricted cells connected by thin ESCRT-III positive cytokinetic bridges. Interestingly, in many of these cells EdU was seen in only one of the two separated nucleoids, indicating that the connecting bridge is not enough to ensure coordinated origin firing in the two daughter cells. Similar results were seen when EdU was visualised using flow cytometry. In control STK positive cells, EdU accumulated in cells with <2N DNA content. By contrast, high levels of EdU were observed in cells expressing Vps4WkB with genome equivalents ranging from 1 to >5, with a peak at ~3.5 genome equivalents **(Fig. 1E, S1F)**. Thus, in stark contrast to the behaviour of cells treated with proteasome inhibitor, Vps4WkB expressing cells continue to undergo repeated rounds of DNA replication following division arrest. In this, *Sulfolobus* cells resemble typical eukaryotes in which origin re-licensing at the end of each cell division cycle requires the proteasome, but not cytokinesis (*31*–*33*).

**Figure 1.**
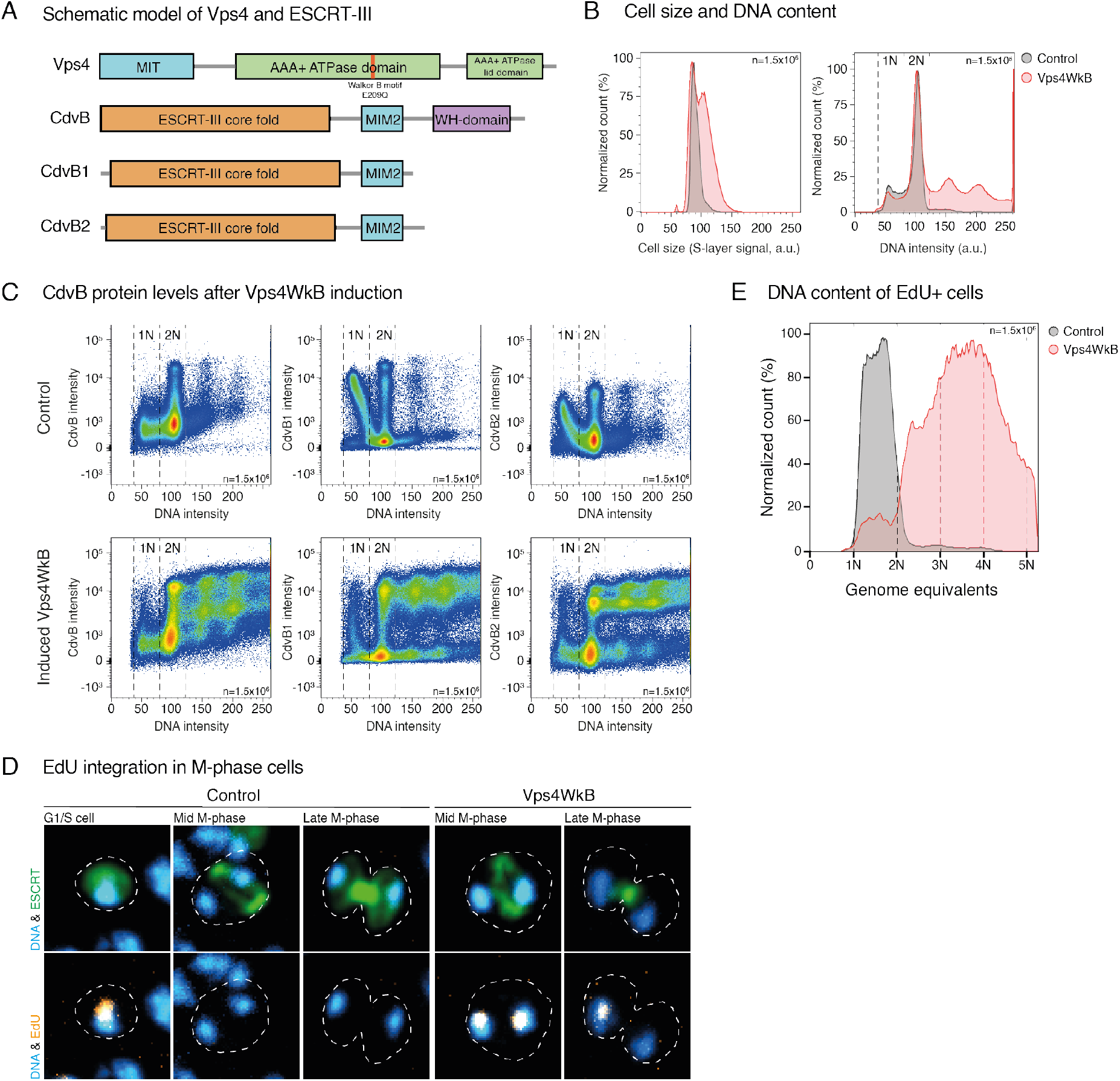
Induction of Vps4WkB causes cell division arrest but not cell cycle arrest. **(A)** Schematic model of Vps4 and ESCRT-III proteins in *S. acidocaldarius* with the approximate location of the Walker B point mutation, E209Q, on Vps4 marked. **(B)** Flow cytograms of control MW001 cells and Vps4WkB cells induced for 8 hours, cell size and 2N+ populations. DNA signal was visualized using DAPI. Cell size approximated from S-layer signal. **(C)** Flow cytograms of control MW001 cells and Vps4WkB cells induced for 8 hours with staining against CdvB, CdvB1 and CdvB2, showing accumulation of ESCRT-III proteins in Vps4WkB induced samples. **(D)** Control and STK+Vps4WkB cells after 4 hours of induction. **(E)** Flow cytometric analysis of cells stained with Edu conjugated to Alexa Fluor 647, showing the population of cells positive for EdU, indicative of DNA synthesis.

To verify that Vps4 activity is required for ESCRT-III mediated cell division, as suggested by this analysis, we upgraded our *Sulfoscope* (*34*) through the addition of a SoRa spinning disk and imaged divisions live in control cells and in cells expressing Vps4WkB that had been pre-labelled with CellMask™ Plasma Membrane Stain (see methods). As previously published (*35*), the rate of cytokinesis (diameter/unit time) in control MW001 cells was roughly constant once cell division had been initiated. By contrast, a short induction of Vps4WkB expression (2 hours) was sufficient to arrest cells at various points in the division process. In almost all cases, the expression of Vps4WkB prevented both ongoing midzone constriction and abscission **(Fig. 2A, S2A)**. Given the well-established role of Vps4 in disassembling ESCRT-III polymers (*26, 36–38*), it seemed likely that this phenotype resulted from an inability of cells to remodel ESCRT-III division rings. In line with this, flow cytometry analysis revealed a significant increase in the percentage of cells positive for CdvB, CdvB1 or CdvB2: from 10% in control cells to 55% after 8 hours of Vps4WkB induction **(Fig. 1C, 2B, S2B-S2C)**. When imaged using immunofluorescence, Vps4WkB expressing cells were associated with an accumulation of division rings at all stages in the division process **(Fig. 2A-2C)**. Importantly, while the constriction of CdvB1 and CdvB2 rings in control cells was associated with a >5-fold increase in cytoplasmic signal for CdvB1 and a 2.5-fold increase for CdvB2 **(Fig. 2D-E)**, indicative of polymer disassembly during cytokinesis (*27*), following Vps4WkB expression the entire pool of CdvB1 and CdvB2 localised to the ring even at late stages of ring constriction **(Fig. 2D-E)**. Importantly, this analysis revealed another clear distinction between the effects of Vps4WkB expression and proteasomal inhibition. While cells treated with Bortezomib are unable to degrade CdvB and arrest with large diameter division rings containing high levels of CdvB, CdvB1 and CdvB2 (*27*), the expression of Vps4WkB blocked rings at every stage of constriction **(Fig. 2C)**. Note, however, that neither Bortezomib treatment nor Vps4WkB appears to impair the ability of cells to assemble CdvB, CdvB1 and CdvB2 rings.

**Figure 2.**
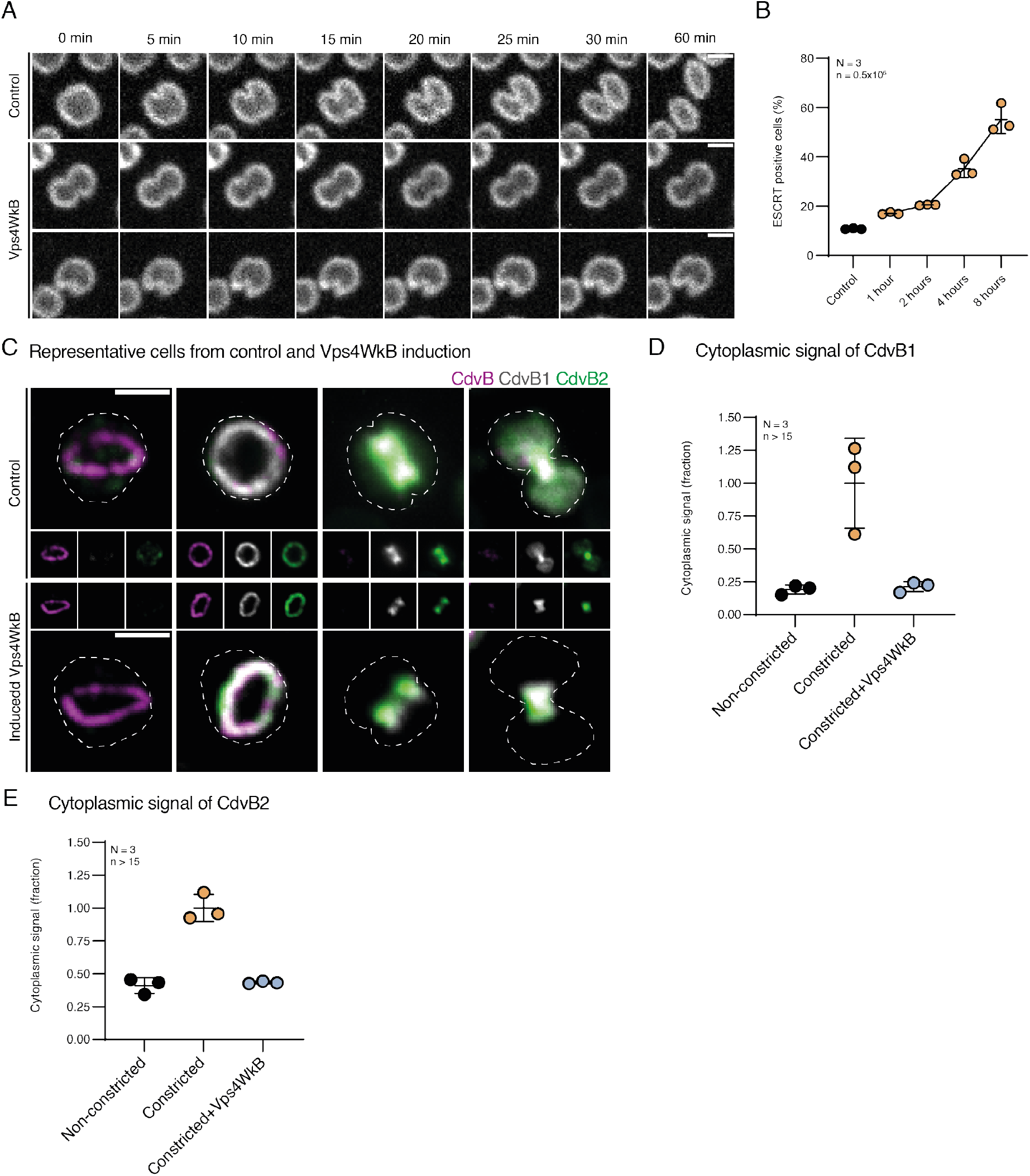
Vps4 is required for cell division. **(A)** Montage of images taken from live imaging of MW001 control and Vps4WkB after 1 hour of induction labelled with CellMask™ Plasma Membrane Stain. Scale bar = 1 μm. **(B)** Quantified flow cytometry data of cells positive for CdvB or CdvB2 in MW001 control or in induced Vps4WkB. Data excludes cells with <2N content. **(C)** Representative immunofluorescent microscopy images of MW001 control and Vps4WkB after 2 hours of induction, larger image showing a composite of the smaller images. Scale bar = 1 μm. Outline of cell shape deduced from S-layer staining. Data from three biological replicates (N = 3) **(D)** Quantification of cytoplasmic signal intensity of CdvB1 in MW001 control and Vps4WkB after 2 hours of induction. **(E)** Quantification of cytoplasmic signal intensity of CdvB2 in MW001 control and Vps4WkB after 2 hours of induction.

Over longer periods of time, the expression of Vps4WkB resulted in even more pronounced division phenotypes - leading to the accumulation of large dumb bell-shaped cells in which the different ESCRT-III proteins occupied a distinct and relatively reproducible position. In these cells, CdvB formed well-defined rings on either side of the cytokinetic bridge, but tended to be excluded from the widest part of the dumbbell and from the narrow portion of the cytokinetic bridge **(Fig. 3A)**. CdvB2 was seen coating the membrane bridge connecting the two daughter cells. In addition, a sub-pool of CdvB2 also colocalised with the CdvB ring as is observed in dividing control cells prior to the onset of CdvB degradation. By contrast, CdvB1 appeared to have a much less well-defined position along the bridge and was seen accumulating in a relatively uniform manner across the entire neck of the dumbbell - extending out from the constricted CdvB2-rich region of the neck beyond the CdvB ring **(Fig. 3A)**. In addition, CdvB1 was observed forming cone-shaped structures in cells with narrow bridges. This same pattern emerged when we averaged the levels of CdvB, CdvB1 and CdvB2 across the cytokinetic bridge of more than 50 cells arrested as dumbbells **(Fig. 3B)**. Again, CdvB localized to the ends of the neck, CdvB2 was preferentially localized to midzone, and CdvB1 spanned the entire length of the bridge. Interestingly, when we repeated the Vps4WkB expression in ΔCdvB1 cells (*34*), CdvB and CdvB2 polymers no longer appeared spatially separated **(Fig. S3A)**. This implies that CdvB1 is required to facilitate the spatial separation of CdvB and CdvB2 polymers in this context. Furthermore, ΔCdvB1 cells expressing Vps4WkB also tended to possess a single CdvB ring rather than two, suggesting that CdvB1 may be required to nucleate a second CdvB ring as cells expressing Vps4WkB continue to grow and cycle without dividing. Note that we were unable to generate ΔCdvB2 cells carrying the Vps4WkB plasmid. The formation of patterned distributions of ESCRT-III proteins over prolonged periods of Vps4WkB expression suggested that different ESCRT-III polymers take up distinct preferred positions in the division ring. Since confocal microscopy limits our ability to observe similar patterns in wildtype cells, we used ~30 nm multi-colour STED microscopy (See Methods) to increase the spatial resolution of our analysis. Strikingly, when visualised by STED, CdvB, CdvB1 and CdvB2 exhibited clear differences in the patterns of accumulation at early stages of division. CdvB was found to localize into a single narrow ring positioned at the centre of control cells **(Fig. 3C)**. Furthermore, CdvB was flanked by two wider accumulations of CdvB1 and CdvB2, which could often be resolved as two distal CdvB1 and two proximal CdvB2 rings **(Fig. 3C-3D, S3B)**. Interestingly, when these rings were observed face on, the diameter of the large CdvB division ring was found to be ~40 nm smaller than the diameters of both CdvB1 and CdvB2 rings **(Fig. S3C)**, implying that CdvB1 and CdvB2 rings may form tighter contacts with the underlaying membrane.

**Figure 3.**
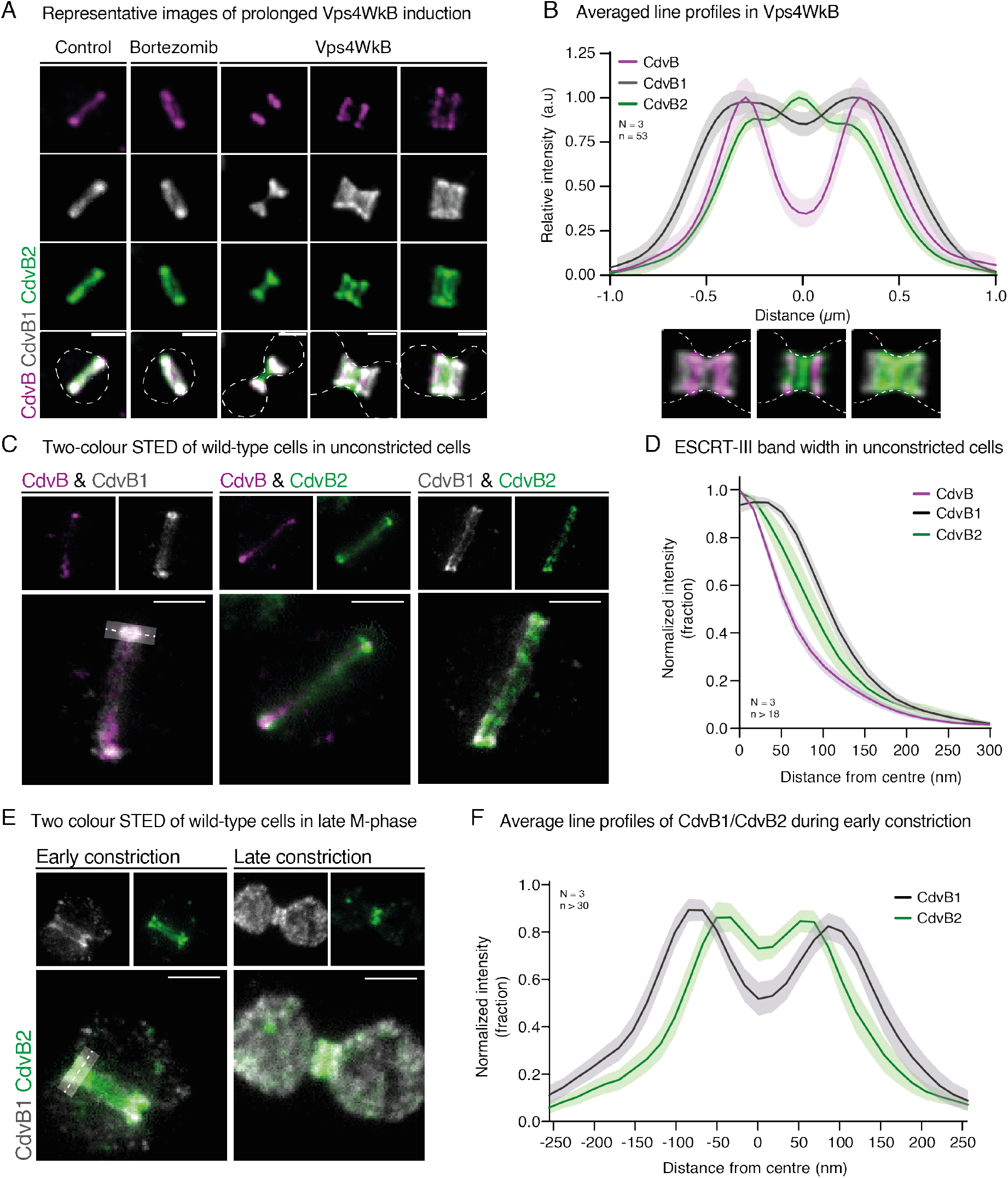
CdvB, CdvB1 and CdvB2 have different relative localization and roles. **(A)** Montage shows prevalent cell morphologies following 8 hours of Vps4WkB overexpression. Outline of cell shape deduced from S-layer staining. Scale bar = 1 μm. N = 3. (B) Quantification of preferred ESCRT-III homologue localization from images like those presented in (A). Data collection used a 2 μm x 25 pixels line profile and was limited to constricted cells expressing all three proteins. Outline of cell shape was defined by S-layer staining. (C) STED microscopy of unconstricted DSM639 wildtype cells. Scale bar = 0.5 μm. Line plot showing the relative localizations of proteins was generated using the average of 5 10-pixel wide, 400 nm long, lines across the ring and fitting a double Lorentzian to the data. N = 3. (D) Line profile quantification of band width from images like those presented in (C). Data collection used a 2 μm x 25 pixels line profile and was limited to constricted cells expressing all three proteins. Outline of cell shape was defined by S-layer staining. (E) STED microscopy of constricted DSM639 wildtype cells. Scale bar = 0.5 μm. Line plot showing relative polymer localizations was generated using a 25-pixel wide, 400 nm long, line across the ring or neck and fitting a double Lorentzian to the data. N = 3. (F) Line profile quantification of band structure from images like those presented in (E). Data collection used a 2 μm x 25 pixels line profile and was limited to constricted cells expressing all three proteins. Outline of cell shape was defined by S-layer staining.

We next examined cells that had begun to constrict following the loss of CdvB. At the early stages of the constriction process, CdvB2 could be seen localizing to a pair of bands in the middle of the cytokinetic furrow flanked by two distal bands of CdvB1 **(Fig. 3E-3F)**. At late stages in the division process, when the wildtype cytokinetic furrow had constricted to a narrow neck, CdvB2 was present in a single linear structure that occupied the central portion of the bridge, whereas CdvB1 was seen throughout the cytoplasm likely as a result of polymer disassembly **(Fig. 3E)** (*27*). Taken together these data suggest a clear order of Vps4-dependent ESCRT-III disassembly from composite division rings: beginning with the removal of CdvB, followed by the disassembly of CdvB1, then CdvB2.

In all systems studied thus far, Vps4 has been shown to be recruited to ESCRT-III polymers via the binding of its MIT domain to C-terminal MIM domains on ESCRT-III proteins as a prelude to polymer disassembly (*6, 24, 39, 40*). To determine how the Vps4-dependent disassembly of each of the three ESCRT-III polymers contributes to division in *S. acidocaldarius* cells, mutant versions of ESCRT-III proteins that lack the corresponding MIM2 domain **(Fig. 4A)** were overexpressed in an attempt to interfere with the stepwise disassembly described above **(Fig. S4A)**. Interestingly, the overexpression of CdvBΔMIM2, led to an accumulation of cells with wide, non-constricted rings containing all ESCRT-III ring proteins, similar to the effect of Bortezomib treatment. This is consistent with the notion that MIM2-dependent removal of CdvB from the ESCRT-III co-polymer ring by Vps4 is a pre-requisite for its subsequent proteasomal degradation as well as constriction of the associated CdvB1 and CdvB2 rings **(Fig. 4B)**. Overexpressed CdvB1ΔMIM2 was found to accumulate in rings **(Fig. S4B),** like CdvBΔMIM2, and was frequently observed in cone-like structures **(Fig. 4C)**. However, CdvB1ΔMIM2 did not appear to interfere with constriction itself. Instead CdvB1ΔMIM2 expression accelerated the rate of constriction measured by live imaging. While control cells constricted at an average rate of 0.11 μm/min (SD = 0.033), CdvB1ΔMIM2 cells displayed an average constriction rate of 0.20 μm/min (SD = 0.061) **(Fig. 4D-4E)**. In contrast, cells lacking CdvB1 were previously shown to take longer to complete division (*34*). Strikingly, CdvB2ΔMIM2 expression had a very different impact from CdvB1ΔMIM2 expression. Cells with high levels of CdvB2ΔMIM2 frequently had rigid **(Movie S1)** spike-like protrusions **(Fig. 4F)**, along with conical bulges and, very occasionally, thin CdvB2ΔMIM2-rich cytokinetic bridges **(Fig. S4C)**. Interestingly, while the expression of CdvBΔMIM2, CdvB1ΔMIM2 and CdvB2ΔMIM2 all led to the accumulation of stable structures, the set of ESCRT-III proteins accumulating in each these structures followed a relatively strict rule **(Fig. S4D)**. CdvBΔMIM2 positive structures contained both CdvB1 and CdvB2, while the conical structures forming in CdvB1ΔMIM2 expressing cells contained CdvB2, but no CdvB. Finally, CdvB2ΔMIM2 positive structures did not contain either of the other proteins **(Fig. S4D)**. These data support a sequential model of ESCRT-III polymer disassembly: CdvB is disassembled first, followed by CdvB1, and then CdvB2. Importantly, while the initial rates of constriction in CdvB2ΔMIM2 cells were slightly reduced as compared to the control **(Fig. 4D, S4E)**, CdvB2ΔMIM2 daughter cells were only very rarely observed to undergo successful abscission **(Fig. 4G)**. Thus, while 70% of control daughter cells separated within 20 minutes from the end of constriction, only 3% of CdvB2ΔMIM2 daughter cells separated within this time frame. These data support the idea that CdvB2 polymers: i) have a preferred curvature compatible with their accumulation in very thin membrane tubes, ii) stabilize membrane tubes, and iii) must be disassembled for abscission to occur. These observations also suggest that a portion of the thin CdvB2-rich spikes are likely to be remnants of CdvB2ΔMIM2 bridges that broke during processing for microscopy. Interestingly, extracellular vesicle formation in *S. acidocaldarius* was also found to be depend on CdvB2, but not CdvB1 **(Fig. S4F)** - in line with CdvB2 playing a crucial role in membrane scission.

**Figure 4.**
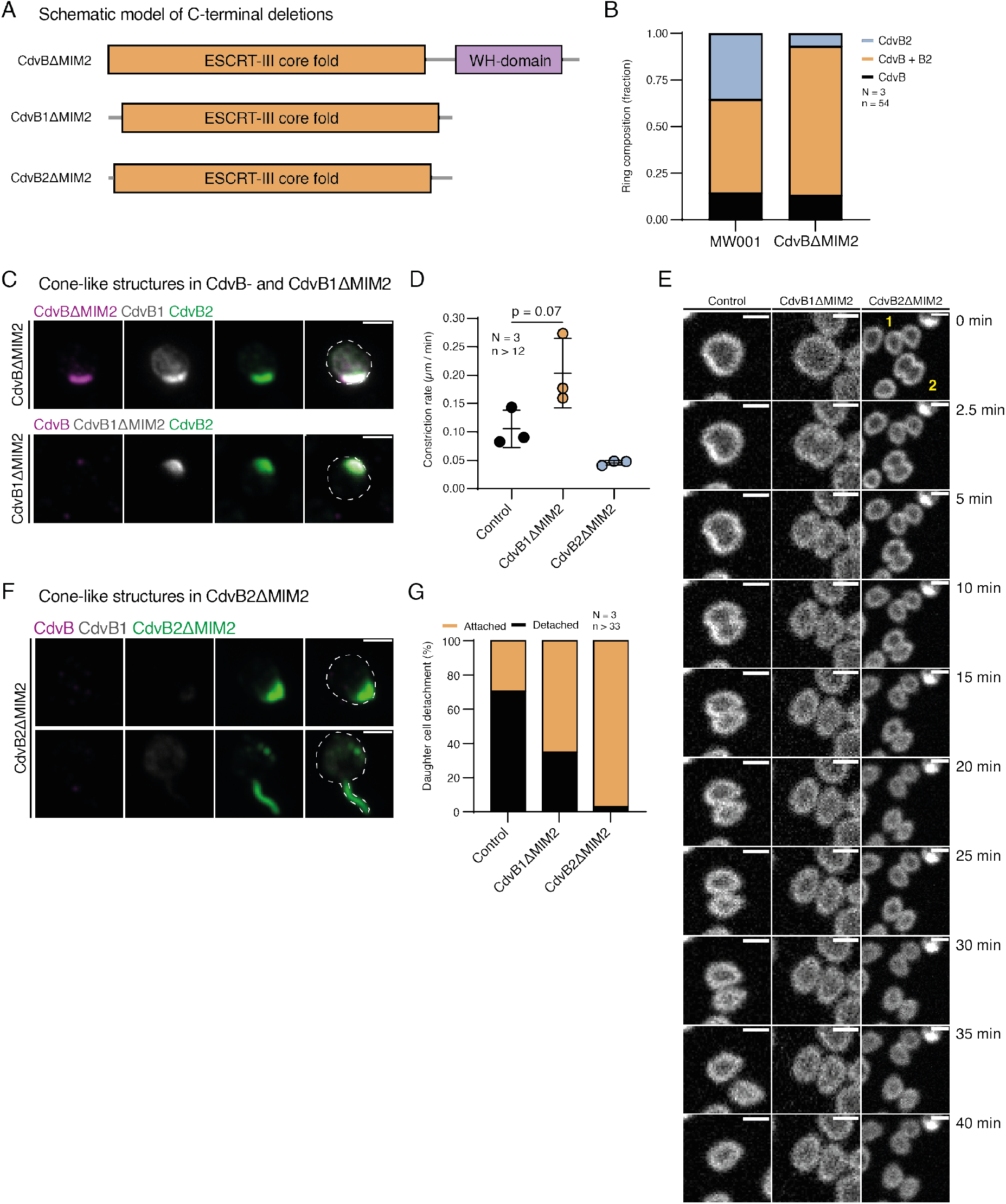
Sequential remodelling of CdvB1 and CdvB2 is required for proper constriction and cell division. **(A)** Schematic of ESCRT-III proteins constructs with MIM2 domain deletions. **(B)** Quantification of microscopy data showing ring composition in MW001 control and CdvBΔMIM2 cells. **(C)** Montages of stills from live imaging of CdvBΔMIM2 and CdvB1ΔMIM2 expressing cells. Scale bar = 1 μm. **(D)** Quantification of constriction speed from live imaging. **(E)** Montages from live imaging of control, CdvB1ΔMIM2 and CdvB2ΔMIM2 expressing cells. Scale bar = 1 μm. **(F)** Quantification of daughter cell separation within 20 minutes of completed constriction, as observed by live imaging of control, CdvB1ΔMIM2 & CdvB2ΔMIM expressing cells. Proportion of successful abscissions quantified. Scale bar = 1 μm). from live microscopy as measured by the proportion of cells in which daughter cells have separated within 20 minutes of end of constriction. Scale bar = 1 μm. **(G)** Montage of immunofluorescence microscopy of CdvB2ΔMIM2 expressing cells showing cones, spikes and long bridges containing CdvB2ΔMIM2 protein. Scale bar = 1 μm.

**Figure 5.**
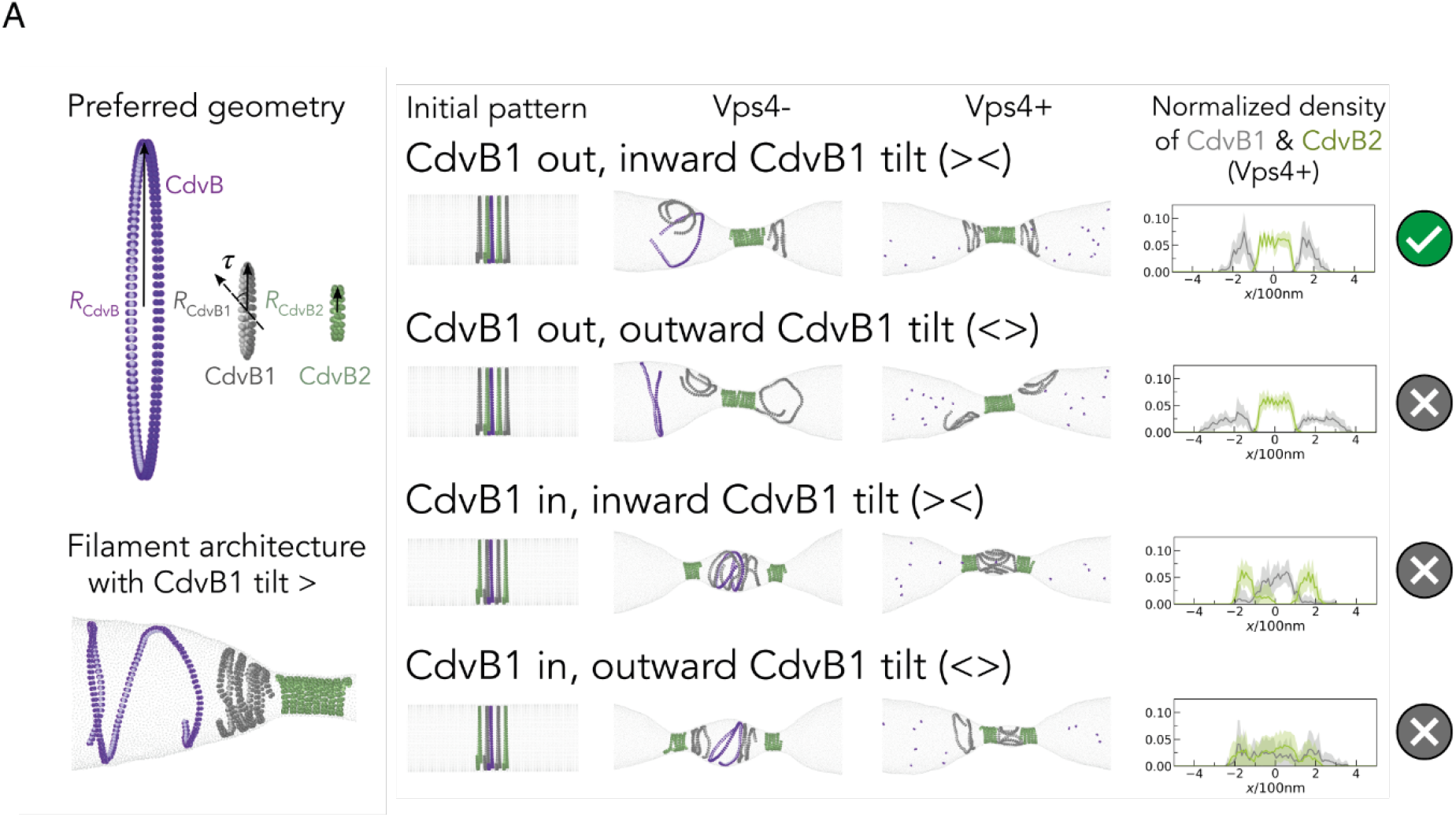
Simulations of membrane constriction in the presence and absence of active Vps4. **(A)** Impact of the recruitment pattern of CdvB1 and CdvB2 on filament constriction and spatial organization without (Vps4-) and in the presence of Vps4 (Vps4+). Left hand panels: the filaments are modelled as short helical strands with 1.05 loops in length with preferred geometries as shown on the left: R_CdvB_ = 170 nm, R_CdvB1_ = 65 nm, R_CdvB2_ = 40 nm, CdvB1 tilt *τ*_*cdvB*1_ = 40° and filament bond stiffness *k_CdvB_* = *k_CdvB2_* = 250 *k_B_T*/*σ*^2^, *k_CdvB1_* = 50 *k_B_T*/*σ*^2^. The membrane is depicted in light grey. Central panels: Initial conditions were defined such that CdvB (purple) forms a single ESCRT-III polymer at the cell centre, which is flanked by CdvB1 (grey) and CdvB2 (green) polymers. Simulation snapshots were then taken as the system was allowed to reach equilibrium as filaments tended towards their preferred curvature. Right: the normalized filament density of CdvB1 and CdvB2 after disassembly of CdvB (Vps4+) was calculated based on data averaged over ten independent simulations - with standard deviations indicated by the shaded areas. The symbol on the far right indicates the chances of different initial patterns of ESCRT-III placement leading to the formation of a single, well-organised constriction.

Taken together, these results imply functionally distinct roles for CdvB, CdvB1 and CdvB2, as the three proteins form homopolymers with distinct localisations and preferential curvatures – something that became evident when these proteins were imaged in cells several hours after the induction of Vps4WkB expression **(Fig. 3A)**. To test the functional significance of this patterned assembly of ESCRT-III polymers we used coarse-grained molecular dynamics simulations (*19, 35*). In this modified computational model, three ESCRT-III polymers representing CdvB, CdvB1 and CdvB2 were pre-assembled together on the membrane near the centre of a tube designed to represent the cell in its pre-division state. Polymers were then allowed to equilibrate and reach their mechanical equilibrium conformation. Since CdvB formed single well-defined rings in non-constricted cells **(Fig. 3C-3D)**, CdvB was modelled as a single polymer that preferentially forms rings with a large radius. By contrast, CdvB2 rings were modelled as helical polymers with a large preferred curvature, in line with observations in wildtype, Vps4WkB and CdvBΔMIM2 cells **(Fig. 3C, 4F, S4C).** Under these assumptions, the parameter space was scanned to determine a set of mechanical properties of CdvB1 that would enable the three polymers to physically separate in simulations **(Fig. 5A).** The spatial separation of polymers occurred more readily when CdvB1 filaments were modelled as being relatively flexible compared to CdvB and CdvB2, with a tilt of the membrane binding interface relative to the helical polymer axis and/or an intermediate preferred curvature between that of CdvB and CdvB2 **(Fig. S5A-S5D)**. The tilt, as defined in Harker-Kirschneck 2019 (*19*), defines the orientation of the polymer’s membrane attraction site relative to its helical axis. When these three properties were combined in simulations, flexible CdvB1 polymers with a tilt of 40° were seen to bind cone shaped membranes as well as the necks of dumbbell-shaped cells, consistent with the pattern of CdvB1 localization observed in Vps4WkB expressing cells **(Fig. 3A)**. Using these parameters, we then tested the importance of precisely positioning the different ESCRT-III polymers on the membrane prior to division, in the presence or absence of Vps4 activity. In these simulations, reproducible formation of a single constriction site required the assembly of two CdvB1 rings placed outside of the two central CdvB2 rings prior to the onset of cytokinesis **(Fig. 5A, S6A, Movie S2A-S2B)**, implying a role for patterned assembly in ESCRT-III dependent division.

## Discussion

Taken together these findings demonstrate an important role for the ordered and pre-patterned assembly of ESCRT-III filaments and their subsequent Vps4-dependent sequential disassembly for cell division in *S. acidocaldarius*. They also suggest a clear model for ESCRT-III dependent cell division. This begins with the assembly of a CdvB ring at the centre of cells that are ready to divide. This single CdvB ring acts as a non-contractile template upon which CdvB1 and CdvB2 can polymerize in a manner that does not depend on Vps4 (*13, 41*).

STED microscopy revealed that CdvB, CdvB1 and CdvB2 formed spatially separated rings in wildtype cells prior to the onset of constriction **(Fig. 3C-3D** and **S3C)**. This suggests that ESCRT-III proteins in *S. acidocaldarius* form layered composite structures, not hetero-polymers like their eukaryotic counterparts Vps2 and Vps24 (*42*). CdvB1 consistently formed two rings flanking CdvB2, while CdvB2 was found either assembled into a single diffuse structure overlying the central CdvB ring or into two flanking rings that are too close to easily resolve using STED. Then, following the onset of cytokinesis, both CdvB1 and CdvB2 signals resolved into two pairs of rings. These data imply that the spatial separation of ESCRT-III polymers may be accentuated as the membrane deforms, something that can also be observed in simulations as polymers seek out regions of suitable curvature **(Fig. 5A)**. Although it is not clear from our analysis exactly how the placement of these different ESCRT-III rings is achieved, these simulations show that a similar spatial separation can be induced by simple differences in the structure and mechanics of the different homopolymers.

During ring constriction, CdvB1 and CdvB2 were found to retain their respective positions: with CdvB1 polymer extending outside of the two central CdvB2 rings. However, once constriction was near complete, CdvB was largely absent and CdvB1 was almost entirely cytoplasmic; leaving cells connected by a thin CdvB2-rich neck. These data suggest that while the majority of the CdvB pool is disassembled and degraded prior to the onset of ring constriction, as previously reported (*22*), the disassembly of CdvB1 begins early in the process of constriction. This is followed, with some delay, by the disassembly of CdvB2. CdvB2 disassembly is likely to be the event that triggers scission in *Sulfolobus acidocaldarius*, since cells expressing CdvB2ΔMIM2 form doublets connected with narrow bridges that fail to separate. These data are in line with data in *Saccharolobus islandicus* (*43*) and with simulations that suggest a role for disassembly in scission (*27, 35*). The importance of CdvB2 for division was also clear from the profound division defect observed in *Sulfolobus* cells lacking the protein; implying that the function of CdvB2 is poorly rescued by the presence of CdvB1 (*34*). Furthermore, the deletion of CdvB2 completely abolished the generation of extracellular vesicles **(Fig. S4F),** while CdvB1 was dispensable for vesicle formation, further supporting a critical role for CdvB2 in membrane scission. The contrast between CdvB1 and CdvB2 function was also clear in MIM2 deletion experiments in which the expression of CdvB1ΔMIM2 was found to accelerate constriction, possibly by increasing the cellular concentration of CdvB1. These data, together with the fact that the loss of CdvB1 leads to longer division times and occasional division failures (*34*), suggest that the flanking CdvB1 polymers facilitate CdvB2 positioning, robust constriction, and CdvB2-dependent scission – as suggested by simulations **(Fig. 5A, S6A-S6B)**.

Having detailed the spatial organisation of ESCRT-III polymers during cytokinesis in *Sulfolobus* cells, it is worth comparing this to what is known about ESCRT-III-dependent abscission in human cells. In the latter case, one or two conical ESCRT-III spirals extend from either side of a stable midbody, leading to the formation of one or two constriction sites at some distance away from the midbody where abscission occurs (*14, 44–46*). By contrast, in *Sulfolobus cells*, the constricting ESCRT-III polymers that drive division appear to take up a symmetrical organisation in which two cones are oriented point to point (> <). In this way the ESCRT-III rings in *Sulfolobus* precisely define a single central site at which membrane remodelling and scission occurs. One of the key reasons for this difference between the archaeal and mammalian systems may be the loss of the CdvB template at the centre of the bridge in *Sulfolobus* cells, which is occluded by the midzone in human cells.

As is the case in human cells, our analysis also shows that cell cycle progression in *Sulfolobus* is uncoupled from ring constriction. Thus, both Vps4WkB and CdvBΔMIM2 expressing cells arrest during division with large rings but undergo extensive endoreplication. This contrasts with the effect of proteasomal inhibition, which arrests both cell division and the cell cycle in *S. acidocaldarius*, implying that the proteasome is needed to degrade an as of yet unidentified protein for origins of replication to be licensed and/or to fire in the following cycle.

In conclusion, this study shows how the patterned assembly and ordered Vps4-dependent disassembly of ESCRT-III copolymers is able to drive ESCRT-III dependent membrane remodelling and membrane scission in a simple *in vivo* model of ESCRT-III/Vps4 function. In this way, our study supports and extends the conclusions of recent *in vitro* studies using the equivalent eukaryotic counterparts, which undergo Vps4-dependent changes in composition (*13*). In the future, it will be important to improve our understanding of the similarities and differences between the process described here and Vps4-dependent ESCRT-III-mediated membrane remodelling in eukaryotes.

## Author contributions

B.B. and F.H. conceived of the study. Initial observations were made by F.H. Cell biology methods and experiments were developed and performed by F.H., A.C. and J.T. with guidance from B.B. Vesicle experiments were carried out by A.A.P. STED methods were developed and performed by T.B., F.M. and R.V. The pipeline of STED image analysis was developed by F.M. and T.B. with support from R.V. Molecular genetics were performed by J.T. and F.H. The physical model was developed by X.J. and L.K. with guidance from A.S. and B.B. Statistical analysis was performed by F.H., F.M., T.B. and A.C.

## Data and materials availability

All data is available in the manuscript or the supplementary materials.

