## Supplemental information for "The patterned assembly and stepwise Vps4-mediated disassembly of composite ESCRT-III polymers drives archaeal cell division"

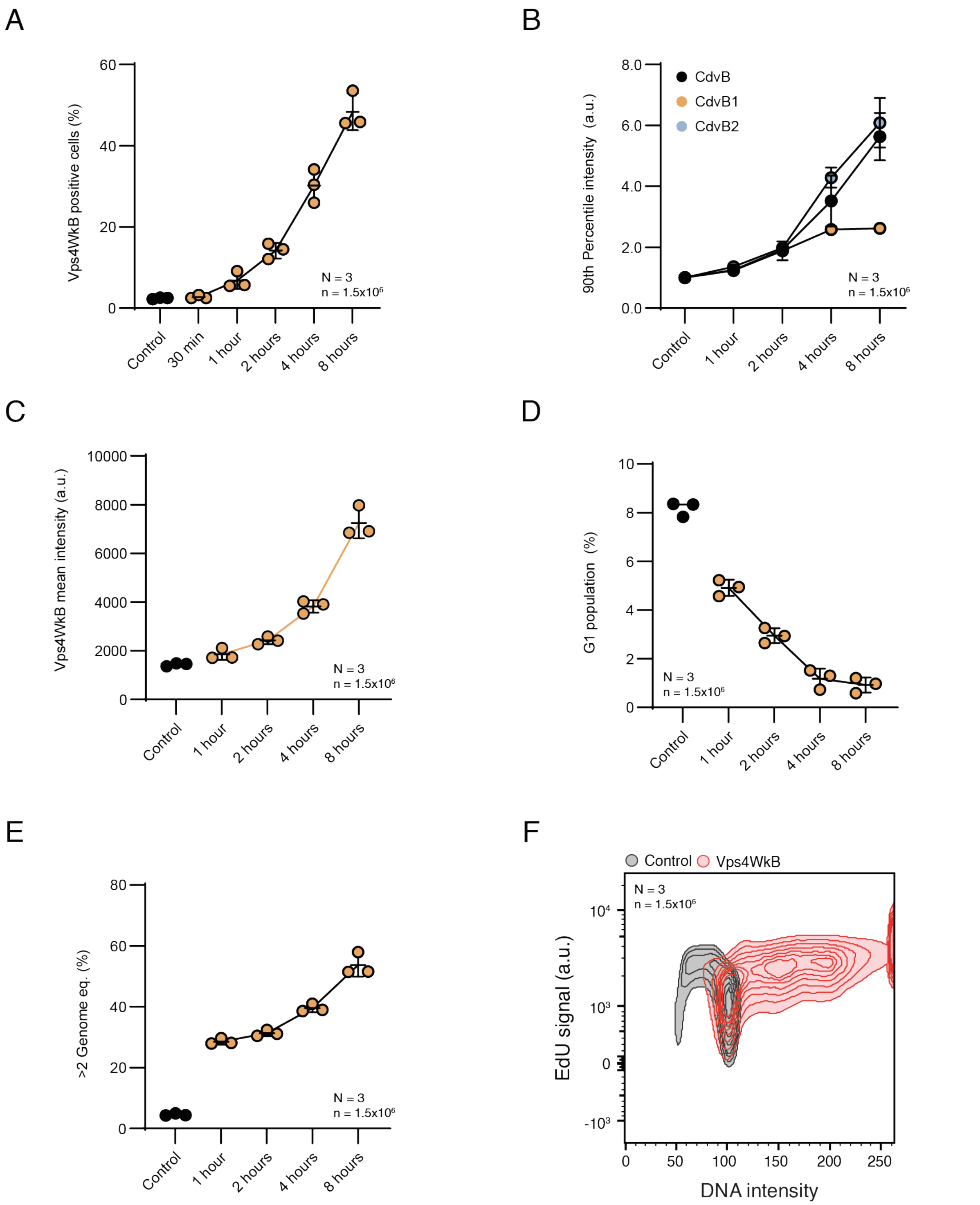


**Figure S1.** Induction of Vps4WkB causes cell division arrest but not cell cycle arrest. **(A)** Quantification of percent of population positive for Vps4WkB signal in flow cytometry. Limit for positive cells determined by control. **(B)** 90% percentile intensity of CdvB, CdvB1 and CdvB2 signal in flow cytometry. **(C)** Mean intensity of Vps4WkB signal in flow cytometry. **(D)** Quantification of G1 population size from flow cytometry. Gating was performed using a DNA vs CdvB2 cytogram. **(E)** Percent of cells with > 2 genome equivalents from flow cytometry in Vps4wkB. **(F)** 2D contour plot showing DNA intensity and EdU signal from flow cytometry. Contour cut off = 10%. N = 3.

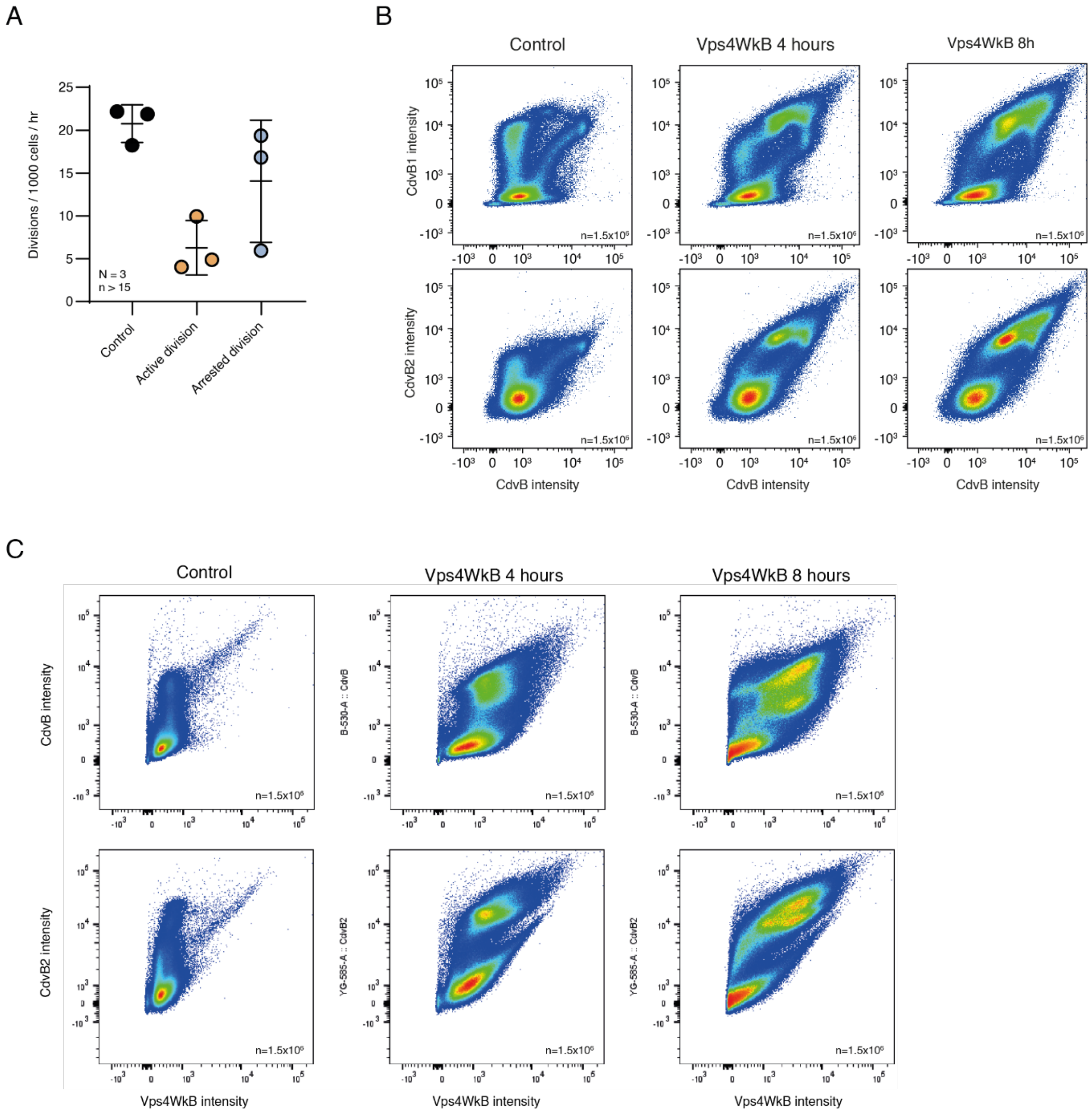


**Figure S2.** Vps4 is required for cell division. **(A)** Quantification of cell divisions from live imaging of MW001 control and Vps4WkB. N = 3 **(B)** Representative cytograms of CdvB protein levels vs CdvB1 and CdvB2 in MW001 control and Vps4WkB. N = 3. **(C)** Representative cytograms of Vps4WkB intensity versus CdvB and CdvB2 in MW001 control and Vps4WkB. N = 3.

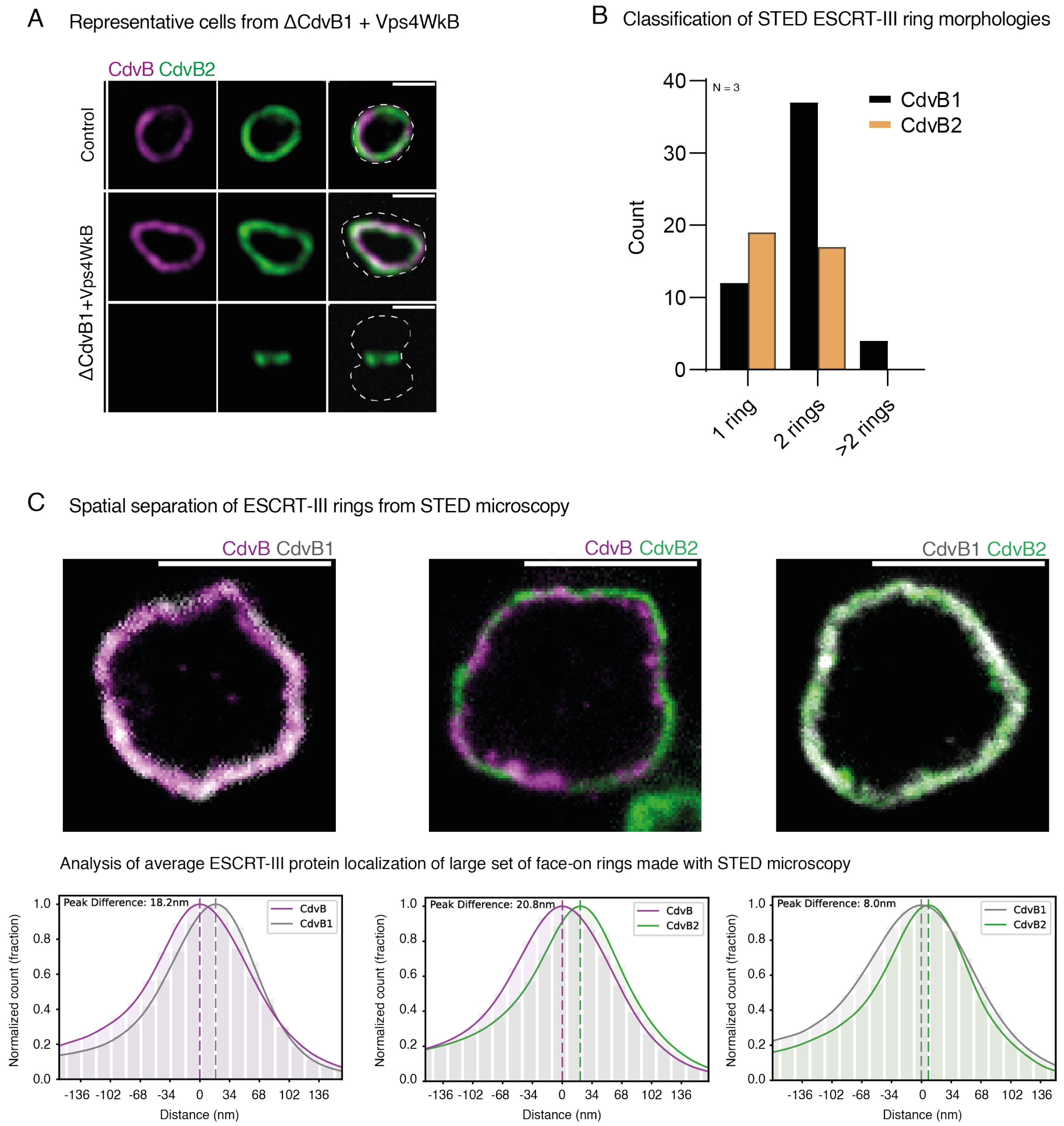


**Figure S3.** CdvB, CdvB1 and CdvB2 have different relative localizations and roles. **(A)** Representative spinning disc microscopy images of MW001 control and ΔCdvB1+Vps4WkB cells. N = 3. **(B)** Classification of ring morphologies from STED microscopy from a large set of images of non-constricted cells retaining a CdvB ring. **(C)** Representative rings when viewed face on by two-colour STED microscopy. Graph shows quantification of averaged ring radii (two per diameter) taken from these data.

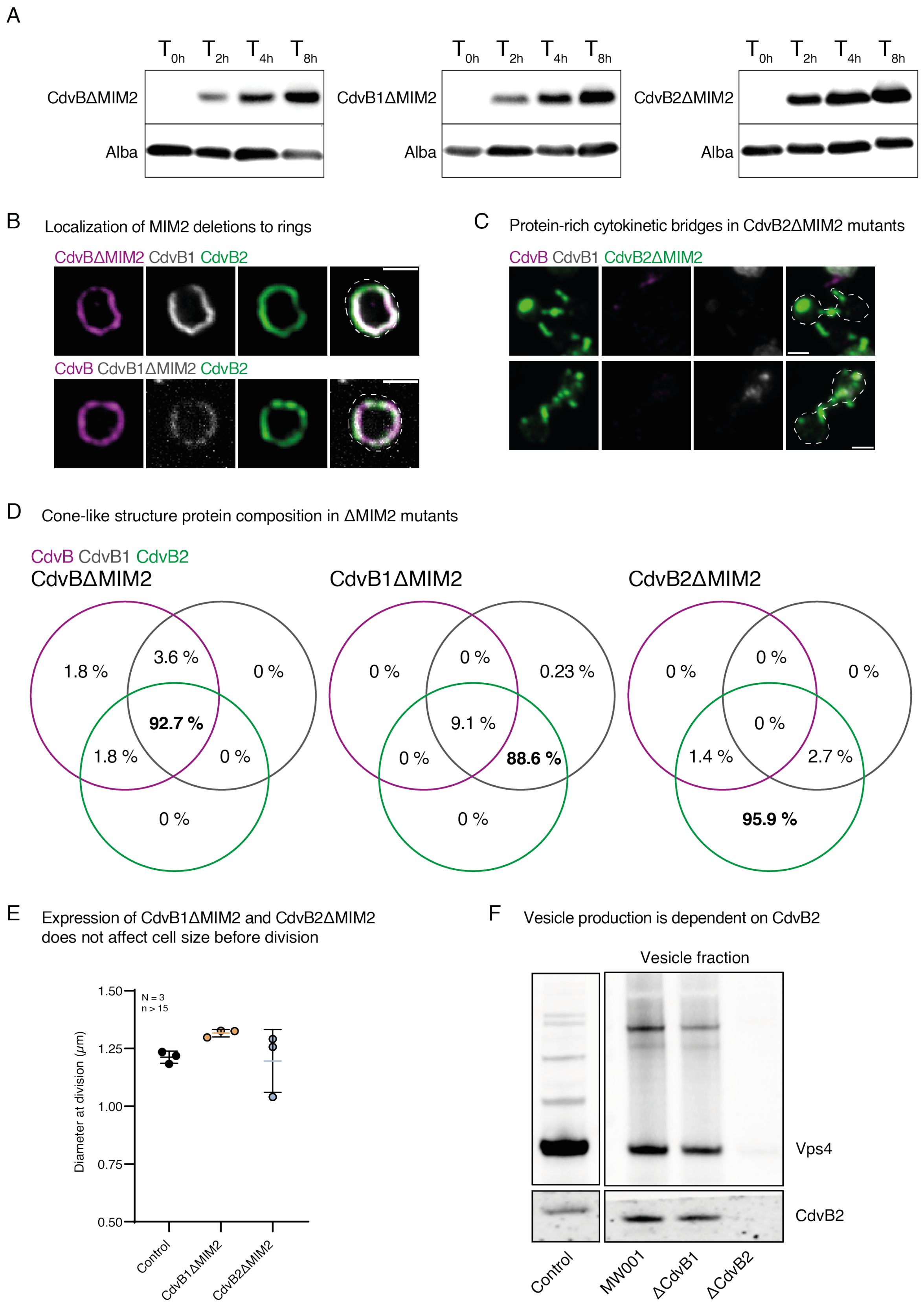


**Figure S4.** Sequential remodelling of CdvB1 and CdvB2 is required for proper constriction and cell division. **(A)** Western blot of ΔMIM2 mutant induction over time, with Alba protein serving as a loading control. N = 3. **(B)** Montages of stills from live imaging of CdvBΔMIM2 and CdvB1ΔMIM2 expressing cells. Scale bar = 1 μm. **(C)** Representative examples of daughter cells connected by narrow bridges with high levels of CdvB2ΔMIM2. N = 3. **(D)** Quantification of CdvB protein presence in cone-like structures present in ΔMIM2 strains. N = 3. **(E)** Quantification of cell size of MW001 control and ΔMIM2 strains from live imaging. N = 3. **(F)** Western blot of MW001 control and vesicles fractions of MW001, ΔCdvB1 and ΔCdvB2, with staining against Vps4 and CdvB2.

Heatmaps exploring the impact of the filament energy on the constriction mechanism

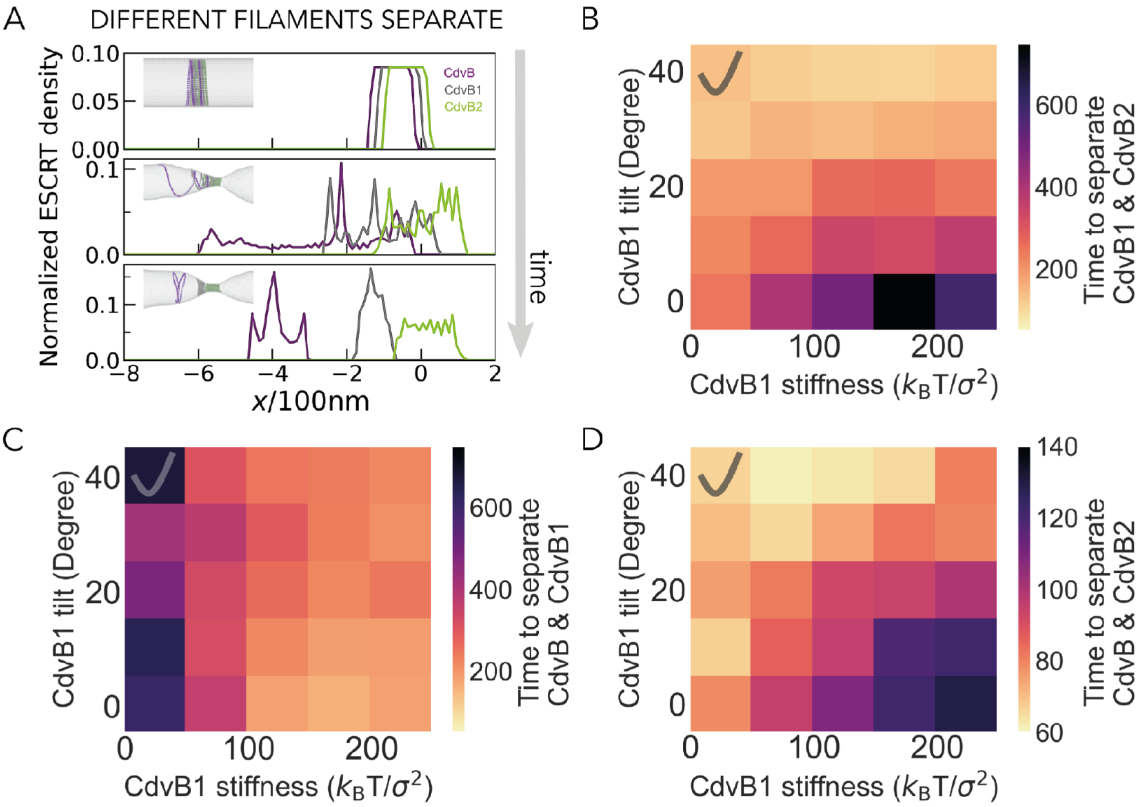


**Figure S5.** Typical snapshots along the spontaneous filament separation trajectory of CdvB, CdvB1, CdvB2 in the absence of Vps4 **(A)** and the time to separate CdvB1 and CdvB2 **(B)**, CdvB and CdvB1 **(C)**, CdvB and CdvB2 **(D)** (in the unit of $500\tau$) as a function of CdvB1 stiffness and tilt. Data is averaged from 5 independent simulations. $k_{CdvB} = k_{CdvB2} = 250\,k_BT/\sigma^2$. In (A), $k_{CdvB1} = 200\,k_BT/\sigma^2$, $\theta_{CdvB1} = 40°$. The biologically relevant parameter regime (CdvB1 soft and tilted) is indicated with a tick label in panels B-D.

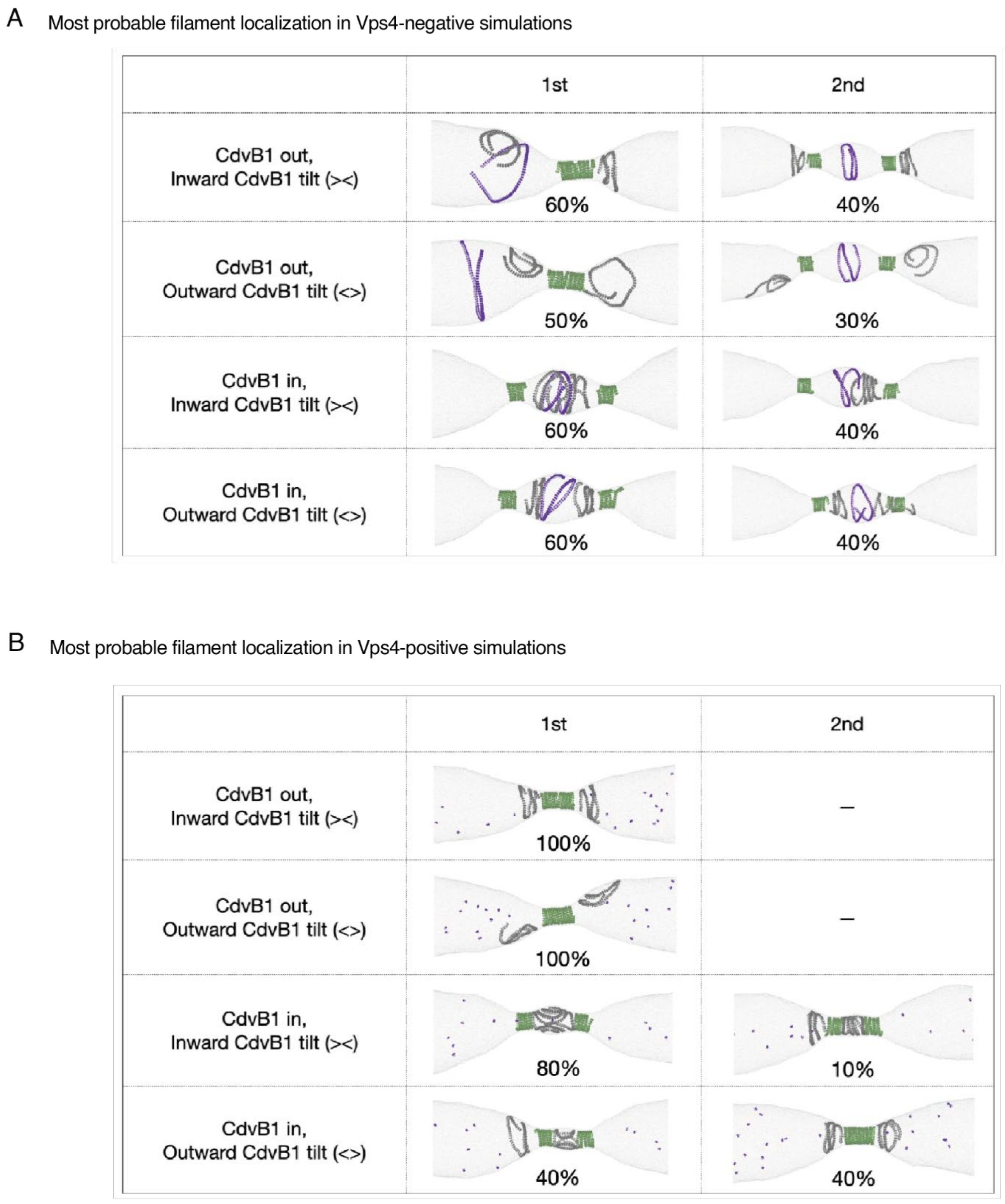


**Figure S6.** Representative snapshots of the first and second probable filament localization in the absence of Vps4 **(A)** and in the absence of Vps4 **(B)**. The most probable configurations are displayed in Fig. 5A.

A Snapshots of a simulation with CdvB1 - CdvB2 - CdvB1 patterning inside a tube

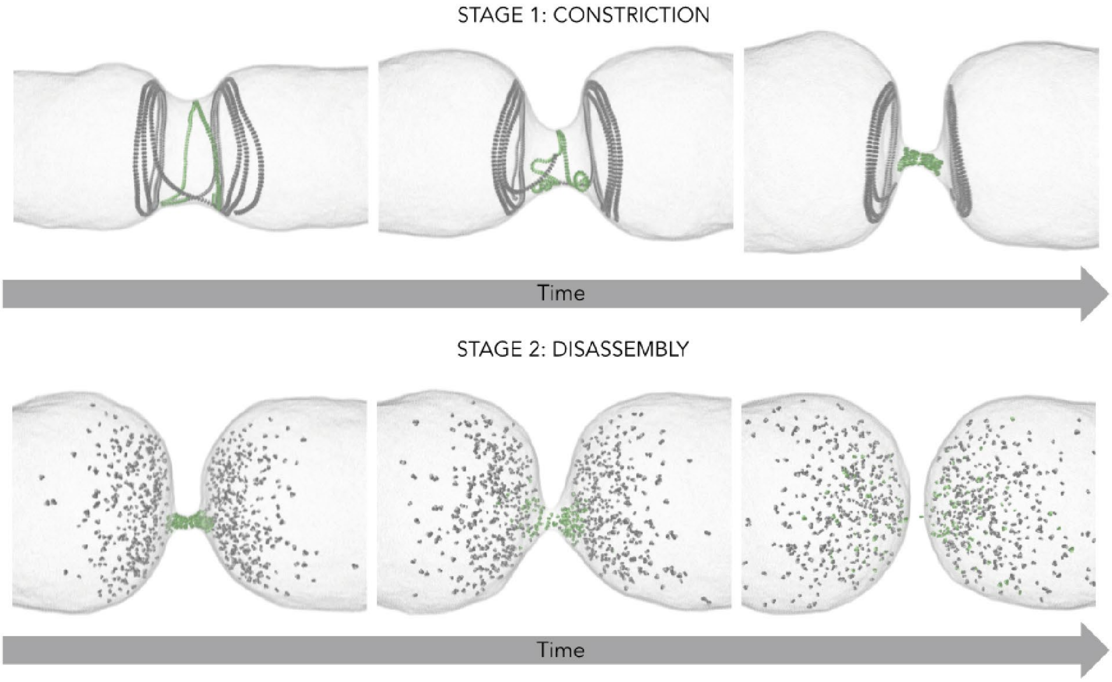


**Figure S7.** Snapshots of a simulation with CdvB1 - CdvB2 - CdvB1 patterning inside a tube with $R_{tube}$ = 27 $\sigma$ The filaments were not supplied with all the energy instantaneously, but instead slowly transformed via transitioning one random subunit within the filament at a time. The initial filament state for both CdvB1 and CdvB2 was R target = R cell, Stiffness = 400k B T/σ 2 and τ = 0°. CdvB1 then transformed to a new state in which R target = 60% R cell, Stiffness = 400k B T/σ 2 and τ = 90° and CdvB2 to R target =1.75 σ, Stiffness = 400 k B T/σ 2 and τ = 0°. This led to CdvB1 forming funnels on each side of CdvB2 and CdvB2 constricting the membrane neck to a tube. Upon disassembly of CdvB1 and then CdvB2 the cell divided, provided the final target radius of CdvB2 was sufficiently small and the membrane was not put under tension.

**Movie S1.** Live cell imaging of *S. acidocaldarius* expressing CdvB2ΔMIM2, displaying rigid membrane protrusions.

**Movie S2.** Coarse-grain molecular dynamics simulations of Vps4+ conditions described in figure 5 and figure 6. **(A)** CdvB1 polymer outside of CdvB2 polymer, inward CdvB1 tilt. **(B)** CdvB1 polymer inside of CdvB2 polymer, inward CdvB1 tilt.

### Methods

#### Cell culturing, fixing and drug treatment

*S. acidocaldarius* DSM 639 (background strain), MW001 (Uracil auxotrophic cloning strain), STK and all plasmid carrying strains were grown in Brock medium pH 2.9 supplemented with 0.1% NZ-amine and 0.2% sucrose at 75°C. MW001 and STK was supplemented with 4 µg/ml uracil where needed. Cells were fixed by stepwise addition of 4°C ethanol to a final concentration of 70% and then stored at 4°C. Proteasome inhibition was performed by treating cells with 10 µM bortezomib (Abcam, ab142123) for 40 minutes. STK strains were supplemented with 200 µM EdU (ThermoFisher, C10639) for 10 minutes.

#### Cloning, transformation and overexpression

The Vps4WkB plasmid was created by amplifying Vps4 (Saci_1372) from *S. acidocaldarius* DSM 639 and generating a point mutation in the Walker B motif at glutamic acid 209 to glutamine and tagged with a 6xHis tag on the C-terminus. CdvBΔMIM2 was generated by amplifying CdvB (Saci_1373) from *S. acidocaldarius* DSM 639 in two blocks, excluding the MIM2 domain (residues 183-193). CdvB1ΔMIM2 and CdvB2ΔMIM2 were generated by amplifying CdvB1 (Saci_0451) and CdvB2 (Saci_1416) from *S. acidocaldarius* DSM 639 up to the point of the MIM2 motif but no further (excluding residues 198-214 and 198-219, respectively). Fragments were cloned into the pSVAaraFX backbone for Vps4WkB and the pSVAaraHA-stop backbone for CdvBΔMIM2, CdvB1ΔMIM2 and CdvB2ΔMIM2 using Gibson Assembly (NEB, E5510S). Plasmids were methylated in vivo prior to transformation into *S. acidocaldarius* by transformation into *E. coli* ER1821 (NEB). Electrocompetent *S. acidocaldarius* MW001 were transformed using electroporation (2000 V, 25 µF, 600 Ω, 1 mm) and selected on Gelrite-Brock plates, following established practices in *Sulfolobus* genetics (*39*).

#### Immunolabelling

Fixed cells were rehydrated in PBSTA (PBS supplemented with 0.2% Tween20 and 3% BSA). Cells were then incubated overnight at 25° C and 500 RPM agitation with PBSTA supplemented with 5% FBS and primary antibodies. Primary antibodies were detected using secondary antibodies for 3 hours at 25°C and 500 RPM agitation. S-layer labelling for cell outlines was performed by incubating cells with Concanavalin A conjugated to Alexa Fluor 488 (ThermoFisher, C11252) or Alexa Fluor 647 (ThermoFisher, C21421) at 50 µg/ml during secondary antibody incubation. DNA was labelled by addition of 2 μM Hoechst (ThermoScientific, 62249) or 3 μM DAPI (ThermoFisher, 62248) after secondary antibody incubation. EdU labelling was performed by a Click-iT kit according to manufacturer instructions (ThermoFisher, C10640). For spinning disc microscopy, Lab-Tek chambered slides (ThermoFisher, 177437PK) were coated with 2% polyethyleneimine (PEI) overnight at RT. Chambers were washed with MilliQ water before stained cell suspension was added and spun down for 1h at 750 RCF. For STED microscopy, 18mm coverslips (Borosilicate #1.5, Marienfeld 0117580) were coated with 2% PEI overnight at RT before washing with MilliQ water. Coverslips were placed in 24-well plates and spun down with labelled cell suspensions for 1 hour at 750 RCF. Coverslips were mounted on microscopy slides using 10 μl of homemade Mowiol mounting media and allowed to cure overnight at room temperature.

Spinning disc microscopy

Cells were imaged in Lab-Tek chambered coverslip using a Nikon Eclipse Ti-2 inverted microscope equipped with a Yokogawa SoRa Scanner unit and Prime 95B sCMOS camera (Photometrics). Images were acquired with a 100x oil immersion objective (Apo TIRF 100x / 1.49, Nikon) using immersion oil (Immersion Oil type F2, Nikon). A total magnification of 280x was achieved using the 2.8x magnification lens in the SoRa unit. Images were acquired with 200 ms exposure time for labelled proteins, and 500 ms exposure time for DNA stains, with laser power set to 20%. Z-axis data was acquired using 10 captures with a 0.22 μm step. Analysis and Z-axis maximum projections were performed using ImageJ and the ImageJ plugin ObjectJ (sils.fnwi.uva.nl/bcb/objectj/).

STED microscopy

STED images were acquired of fixed samples using an Abberior Expert Line STED microscope with a 100x oil immersion objective (Olympus Objective UPlanSApo 100x/1.40 Oil). Prior to imaging the confocal and STED lasers were aligned using a 0.1 μm TetraSpeck™ (Invitrogen, T7279) bead sample. Additionally, by switching secondary antibodies, we verified that none of the relative positions of the polymers were affected by the used fluorophores.

Fluorophores are subjected to photobleaching, limiting the cycles of excitation and de-excitation, before they transition into a non-fluorescent state. To minimise this photobleaching, adaptive illumination strategies RESCue (Reduction of State transition Cycles) and DyMIN (Dynamic Intensity MINimum) were utilised (39, 40). In RESCue-mode, intensity thresholds are set at specific percentages of the dwell time. If these thresholds are not reached, the lasers will be switched off for the remainder of the dwell time. As a result, imaging only occurs if a structure is present. In DyMIN-mode, an intensity threshold was set on the confocal image, creating a low-resolution structure localisation mask. A second, more precise structure localisation was determined by imaging within the masked region using a low-intensity STED beam. For the final image, a high-intensity STED beam was used to achieve the best attainable resolution.

All STED images were acquired with a pixel size of 17nm and a 0.8 a.u. pinhole. For excitation, we used 40 MHz pulsed lasers with a wavelength of 561 nm (200 µW at laser head) and 640 nm (1 mW at laser head). The STED depletion laser (also 40 MHz pulsed) has a head intensity of 3.2 W at 775 nm. We applied gated detection, with a window of 13 ns in the 640 channel, and 14.3 ns in the 561 channel. The applied laser powers and dwell times (line steps multiplied with pixel dwell time) were optimised for the different samples as shown in **Table S1**. For the detection of the emission into the 640 channel (640 nm laser excitation), the detection filter was set to 697 ± 56 nm. For the detection of the 560 channel (561 nm laser excitation), the detection filter was set to 603 ± 27nm.

Control samples with secondary antibodies on slides were prepared to estimate the achieved resolution. First, coverslips were coated with poly-L-Lysine (PLL) for 2 hours and washed with MilliQ afterwards. A droplet of 10 µL of secondary antibody, Abberior STAR 580 (Abberior, ST580-1002-500ug, lot# 90619CWF-3) or Abberior STAR 635 (Abberior, ST635-1002-500UG, lot# 18052018-Hp), was pipetted on a piece of parafilm in a humid chamber. The PLL-coated coverslip was placed on top of the droplet and was incubated at room temperature for 1 hour. After incubation, 6 µL of homemade Mowiol was pipetted on a glass slide, and the coverslip with secondary antibody was put on top and left to dry overnight at room temperature, protected from light. For the imaging of these control samples, settings matching all STED measurements were used. For the analysis, ImageJ was used to draw line profiles and measure the FWHM as a measure for the resolution, resulting in an estimated resolution of 29.6 ± 5.62 nm (n = 33, ±1 STD).

**Table S1.** Overview of the STED DyMIN settings for the ESCRT-III proteins CdvB, CdvB1 and CdvB2 in either the 640nm channel (protein labelled with Abberior STAR 635) or the 561nm channel (protein labelled with Abberior STAR 580). Laser intensities are indicated in percentages of the laser heads, where the 640 nm laser power is 1mW and the 561nm laser has 200µW power. The dwell time of a single line was consistently 13.5 µs.

| 640nm channel | 561nm channel | Line steps | | | Intensity Confocal laser (%) | | | Intensity STED doughnut (%) | |
|---|---|---|---|---|---|---|---|---|---|
| | | Confocal | Low int. STED | High int. STED | Confocal | Low int. STED | High int. STED | Low int. STED | High int. STED |
| CdvB | | 5 | 6 | 19 | 0.4 | 0.5 | 1.1 | 4.6 | 45 |
| | CdvB | 2 | 8 | 8 | 7.8 | 13 | 20 | 15 | 90 |
| CdvB1 | | 3 | 5 | 18 | 0.2 | 0.45 | 1.1 | 3.6 | 45 |
| | CdvB1 | 1 | 4 | 5 | 3 | 4 | 15 | 12 | 90 |
| CdvB2 | | 3 | 5 | 18 | 0.2 | 0.45 | 1.1 | 3.6 | 45 |
| | CdvB2 | 2 | 10 | 8 | 17 | 20 | 25 | 15 | 90 |

### Flow cytometry

DNA was labelled with 1.25 nM DAPI for flow cytometry. Flow cytometry analysis was performed on BD-Biosciences Fortessa. Lasers wavelengths and filters used were as followed 355 nm, 488 nm, 561 nm, 633 nm, with filters respectively: 450/50 UV, 525/50 Blue, 582/15 YG, 710/50 Red. Side Scatter and Forward scatter was recorded. Analysis was performed using FlowJo v10.8.1.

### Western blotting

Cells were lysed in Laemmli buffer, followed by 4 cycles of sonication (30/30 on/off) at low power settings on a Bioruptor Plus Sonication System (Diagenode, B01020001). Before running samples on gel, samples were incubated at 99° C for 10 minutes. 10 µL of samples were run on NuPAGE 4-12% Bis-Tris gels (Invitrogen) at 180 V using MES running buffer. Proteins were transferred to a nitrocellulose membrane and blocked with PBST + 5% milk for 1 hour, after which membrane stained with primary antibodies in PBST + 5% milk and 5% FBS overnight at 4° C. Membrane was washed three times in PBST for 5 minutes and then incubated with PBST + 5% milk and secondary antibodies for 2 hours. Membrane was recorded using a Bio-Rad ChemiDoc system.

Primary antibodies

| Antibody (animal) | Supplier |
|---|---|
| Anti-CdvB (rabbit) | Lab generated |
| Anti-CdvB1 (chicken) | Lab generated |
| Anti-CdvB2 (guinea pig) | Lab generated |
| Anti-Vps4 (rabbit) | Lab generated |
| Anti-HA (mouse) | ThermoFisher, 26183 |
| Anti-His (mouse) | Abcam, ab18184 |

Secondary antibodies

| ThermoFisher | Anti-Rabbit | Anti-Chicken | Anti-Guinea pig | Anti-Mouse |
|---|---|---|---|---|
| Alexa Fluor 405 | A48254 | A48260 | | A31553 |
| Alexa Fluor 488 | A11034 | A11039 | A11073 | A10631 |
| Alexa Fluor 546 | A11035 | A11040 | A11074 | A11030 |
| Alexa Fluor 647 | A21245 | A21449 | A21450 | A21235 |

STED secondary antibodies

| | |
|---|---|
| Abberior Star 580 goat anti-rabbit IgG | ST580-1002-500UG |
| Abberior Star 635 goat anti-rabbit IgG | ST635-1002-500UG |
| Abberior Star 580 goat anti-chicken IgY | ST580-1005-500UG |
| Abberior Star 635P goat anti-chicken IgY | ST635-1005-500UG |
| Abberior Star 580 goat anti-guinea pig IgG | ST580-1006-500UG |
| Abberior Star 635 goat anti-guinea pig IgG | ST635-1006-500UG |

Live imaging

Live cell imaging was performed using the Sulfoscope microscopy set up as described in (*29*), with minor modifications. Briefly, 25 mm coverslips were washed with EtOH and $H_20$ then assembled into commercial Attofluor chambers and covered with 300 µl BNS. Chambers were incubated at 75˚C until the media had dried, then washed thoroughly with BNS and placed into the Sulfoscope to equilibrate to 75˚C. For visualisation of S. acidocaldarius membranes, cells at OD600nm 0.15 - 0.3 were supplemented with 0.75 µg/ml CellMask Deep Red Plasma Membrane Stain (ThermoFischer) immediately prior to imaging. For imaging, 400 µl cells were loaded into the Attofluor chamber and immobilised using heated, semi-solid BNS pads (0.6% Gelrite, 0.5x BNS (pH ~5) and a final concentration of 20 mM $CaCl_2$). Immobilisation pads were prepared by cutting 7mm length half-moon shapes from semi-solid BNS media (15ml/plate), placed onto 13mm circular coverslips and incubated at 75˚C for 5-10 min in a bead bath until the pads had slightly dried and the edges of the pad curved downwards. Pre-heated pads were then placed in the chamber such that the edge of the pad was in the centre of the chamber. Cells found at the border of the immobilisation pad were imaged as these cells are only subject to reduced diffusion resulting in a moderate level of immobilisation rather than directly in contact with the pad, limiting the mechanical stress they are under. Cells were allowed to settle for 5-10 min before acquisition on a Nikon Eclipse Ti-2 inverted microscope equipped with a Yokogawa SoRa Scanner unit and Prime 95B sCMOS camera (Photometrics). Images were acquired with a 60x oil immersion objective (Plan Apo 60x/1.45, Nikon) using a custom formulated immersion oil for high temperature imaging (maximum refractive index matching at 70˚C, n = 1.515 ± 0.0005, Cargille Laboratories). A total magnification of 168x was achieved using the 2.8x magnification lens in the SoRa unit. Images were acquired with 15ms exposure time and 10% laser power at intervals of 15s (one colour imaging) or 20s (dual colour imaging) for 2-3hrs. After acquisition, the ImageJ plugin StackReg (*40*) was used to correct XY drift.

**Supplementary Text**

Computational details

The ESCRT-III filaments were initialized as five short polymers placed successively along the inner surface of a long, tubular, deformable membrane **(Fig. 5A)**. To model the filaments, we used the ESCRT-III model we developed in Harker-Kirschneck 2019 (*14*), in which the filament consists of three beaded monomers that are bonded to their neighbouring monomers via nine harmonic bonds. The bond strength determines the stiffness of the polymer, whereas the exact ratio of the bond lengths give the filament its chiral curvature. The filament can constrict or expand its target radius by modifying its bond lengths.

Here each filament consisted of 1.05 helical loops (82 monomers). Two filament recruitment patterns were investigated as the initial configuration of the molecular dynamics (MD) simulations: CdvB1-CdvB2-CdvB-CdvB2-CdvB1 (termed “CdvB1 out”), CdvB2-CdvB1-CdvB-CdvB1-CdvB2 (termed “CdvB1 in”). The target radii of CdvB (purple), CdvB2 (green), CdvB1 (grey) were set as $R_{CdvB} = 17\sigma$, $R_{CdvB2} = 4.0\sigma$, $\frac{1}{R_{CdvB1}} = \left(\frac{1}{R_{CdvB}} + \frac{1}{R_{CdvB2}}\right)/2$ (i.e., $R_{CdvB1} = 6.5\sigma$), where $\sigma$ is the MD unit of length and corresponds to roughly 10 nm. Filament bond stiffness was set as $k_{CdvB} = k_{CdvB2} = 250\ k_B T/\sigma^2$, $k_{CdvB1} = 50 k_B T/\sigma^2$. CdvB1 was given a tilt $\tau_{CdvB1} = 40°$, and we investigated whether the outward (< >) or inward (> <) conical direction at filament recruitment had an impact on the filament separation and constriction. The membrane was modelled with the one-particle-thick model developed by Yuan 2010 (*41*). The membrane tube consisted of 5000 beads and had an initial radius of $R_{tube} = 18\sigma$. The tube length in x direction was set to $l_{tube} = 360\ \sigma$. The size of the coarse-grained beads was $r_{membrane} = r_{subunit} = 0.5\ \sigma$. Short-ranged 12-6 Lennard-Jones (LJ) interactions were applied between the two bottom beads of the filament's three-beaded subunits and the membrane beads:

$$E_{ij} = 4\epsilon\left(\left(\sigma/r_{ij}\right)^{12} - \left(\sigma/r_{ij}\right)^{6}\right) - E_c\ , r_{ij} < r_{cut}.$$

The interaction strength was set to $\varepsilon_{LJ} = 3.0\ k_BT$ and the cut-off distance to $r_{cut} = 1.3 \cdot r_{min}$, where $r_{min} = 2^{\frac{1}{6}}\sigma$ is the inter-particle contact distance and $E_c$ is the energy at cut-off distance. Volume exclusions was applied between different filament subunits and between the membrane and the top beads of the filament. The volume exclusion interactions were treated as LJ interactions truncated at $r_{min}$ and shifted to zero, with the interaction strength set as to $\varepsilon_{volex} = 2.0\ k_BT$. Periodic boundary conditions were applied in three dimensions. We performed molecular dynamics simulations using the molecular dynamics package LAMMPS (*42*) and evolved the equations of motion in the $N_{particles}V_{box}E_{system}$ ensemble coupled to a Langevin thermostat. The Langevin thermostat was applied at every step with the temperature set to 1 and the damping coefficient set to 1 $t_0$ units, where $t_0$ is the MD unit of time. The box size was kept constant. Initially the system was equilibrated for t=100$t_0$, with all three filament types having the same target radius $R_{CdvB} = R_{CdvB1} = R_{CdvB2} = 17\ \sigma$, where $t_0$ is the MD unit of time. This allowed the membrane to attach to the filament loops from the outside. Then CdvB remained at $R_{CdvB} = 17\ \sigma$, while CdvB1 and CdvB2 reduced their target radius to $R_{CdvB1} = 6.5\ \sigma$ and CdvB2 to $R_{CdvB2} = 4.0\ \sigma$ respectively. We then let the system to evolve for $t = 2\times10^5\ t_0$, until the formation of a stable filament distribution formed. In Vps4+ simulations, we disassembled the CdvB filament by severing its internal bonds and turned the interaction between bottom beads and membrane beads from short-ranged LJ interactions to volume exclusions immediately after the initial equilibration. We carried out ten independent simulations for each setup and the configurations of the last snapshots of the trajectories were used to calculate the normalized filament density **(Fig. 5A, S6A-S6B)**. We visualize our simulation results using OVITO (http://ovito.org).

To see if this newly found filament alignment (CdvB1-CdvB2-CdvB1 with CdvB1 tilted inwards) can cause cells to divide, we placed two opposing CdvB1 filaments (each three loops long) around a single looped CdvB2 filament inside a membrane tube with $R_{tube} = 27\ \sigma$. In this simulation filament curvature is changed progressively, by picking random monomers within the filament and constricting them, instead of releasing all the filament energy at once **(Fig. S7A)**. As shown in Harker-Kirschneck et al. 2022 with a single filament, this dynamical protocol led to the most reliable and symmetric division and came remarkably close to matching the kinetics of ring constriction measured from experimental data (*30*). In this new simulation setup with multiple ESCRT-III filaments, the cubical simulation box had a length of 400 $\sigma$ and CdvB1 was given a large tilt $\tau_{CdvB1} = 90°$. The large tilt allowed CdvB1 to decrease the diameter of the membrane neck by providing a surface for the membrane to glide over and into the neck. The filament stiffness was set to $k_{CdvB1} = k_{CdvB2} = 400\ k_B T/\sigma^2$. After equilibrating the membrane on its own for 200 $t_0$, we started by randomly transitioning the CdvB1 filaments from the initial state ($R_{CdvB1}$= $R_{tube}$, $\tau_{CdvB1}$ = 0°) to its tilted and constricted final state ($R_{CdvB1}$ = 60% $R_{tube}$, $\tau_{CdvB1}$ = 90°). Every 100 simulation steps one subunit in each CdvB1 filament got transformed, while the CdvB2 filament remained in its initial state ($R_{CdvB2}$ = $R_{tube}$). Then the CdvB2 filament in the middle was transitioned by transforming one random subunit within the filament from a large to a small target radius every 10000 steps, until the entire filament was transitioned. This transition was much slower, as the target radius of CdvB2 has to decrease dramatically from $R_{CdvB2} = R_{tube}$ to $R_{CdvB2} = 1.75\sigma$ – by about 93.5%. After constriction was completed, we disassembled the filaments by severing the bonds between random subunits within the filament – first CdvB1 and then CdvB2, which led to division. We found that scission only happens reliably if i) the target radius is very small, $R_{target} \leq 2\sigma$, and ii) we use the NPH ensemble instead of NVE as an integrator for the membrane, as that does not put the membrane under tension, but instead lets the membrane area shrink/increase as required (by adjusting the simulation box size). The pressure was kept at 0 to create a membrane under low tension. This supports the finding of Lafaurie 2013, who show that membrane tubes under tension don't divide (*43*).

When simulating filament separation, we initially placed the filaments on the membrane as three interlaced strands, consisting of two full helical loops **(Fig. S5A)**. Each filament strand was modelled by our three-bead-monomer ESCRT-III model (*14*) and consisted of 156 monomers. The membrane tube consisted of 30000 beads. The initial radius of the membrane tube was $R_{tube} = 18\sigma$, with length in x direction $l_{tube} = 216\sigma$. Initially the system was equilibrated for t = $100 t_0$ with CdvB, CdvB1 and CdvB2 all at the same target radius $R_{CdvB} = R_{CdvB1} = R_{CdvB2} = 17\sigma$, to ensure that all three filaments attached to the interior of the membrane tube. We then activated CdvB1 and CdvB2 by resetting their target radii, and allowed the system to evolve until the filaments were fully separated (typically $10^5$ to $10^6$ $t_0$). Time step $dt = 0.01 t_0$ was used for all MD simulations.

<u>Characterization of filament separation time (Fig. S7)</u>

Firstly, we computed the overlap function (OF) by integrating the overlap of the normalized bead distribution along the x-axis of the two filaments of interest. OF = 0 means fully separated and OF = 1 means fully overlapped. The OF was averaged over five independent simulations. The filament separation time is defined as the first time when the averaged OF is below a certain threshold value. The threshold value was computed from the average of the OF over the last 100 frames.
